# Myelin maturation failure and oligodendrocyte precursor loss underlie default mode network disruption in a humanized APP knock-in model of Alzheimer’s disease

**DOI:** 10.64898/2026.08.12.744193

**Authors:** Esmin Unaran, Olamide Adebiyi, Medha R. Krishnan, Timothy J. Bussey, Lisa M. Saksida, Marco A. Prado, Vania F. Prado, Ravi S. Menon

## Abstract

Alzheimer’s disease disrupts large-scale brain networks before cognitive symptoms emerge, yet the mechanisms underlying this progression remain unclear. In humans, default mode network (DMN) connectivity follows a biphasic trajectory, with a transition from hypersynchrony to hyposynchrony associated with increasing amyloid burden and cognitive decline. Using a mouse model of amyloidopathy that recapitulates this trajectory, we show that amyloid-β accumulation arrests normal age-dependent myelin maturation, followed by progressive myelin loss and oligodendroglial dysfunction. In healthy animals, regional myelin coverage predicts DMN strength, revealing a coupling that is disrupted by amyloidopathy. Notably, DMN hypersynchrony emerges before substantial amyloid accumulation or overt myelin loss, whereas subsequent oligodendroglial dysfunction and myelin loss coincide with DMN disintegration and cognitive deficits. These findings identify disruption of myelin–network coupling as a potential mechanism underlying the biphasic evolution of network dysfunction in Alzheimer’s disease and suggest that restoring oligodendroglial function may provide cognitive benefits beyond those of amyloid-targeting strategies.

## Introduction

Alzheimer’s disease (AD) pathogenesis is characterized by a complex interplay of neuropathological processes involving both neuronal and non-neuronal cell types^1,2^. Classically defined by the accumulation of extracellular amyloid-beta (Aβ) plaques and intracellular neurofibrillary tangles composed of hyperphosphorylated tau, AD is accompanied by widespread synaptic dysfunction and progressive neurodegeneration throughout distributed brain regions^3^.

Despite decades of research, therapeutic strategies targeting these hallmark pathologies have yielded limited clinical benefit to memory and other cognitive symptoms that are meaningful to patients. Furthermore, current FDA-approved amyloid-clearing interventions carry significant risks, including amyloid-related imaging abnormalities (ARIA) and brain volume loss, particularly in APOE4 carriers^4–7^. These limitations have shifted attention toward the broader neuropathological landscape of disease progression in AD, including the contributions of non-neuronal cell types and white matter dysfunction^8,9^.

Resting-state functional magnetic resonance imaging (rs-fMRI) studies have demonstrated that large-scale brain network alterations emerge early in AD, preceding overt neurodegeneration and correlating with cognitive decline^10–14^. Among affected networks, the default mode network (DMN) including the cingulate cortex, hippocampus, and sensory regions is among the earliest and most consistently affected^15^. The DMN was initially defined as a set of co-varying brain regions exhibiting high baseline metabolic activity during rest that are systematically suppressed during goal-directed behavior, revealing an organized “default” state of brain function^16^. Over the past two decades, large-scale meta-analytic evidence has established the DMN as a core, actively engaged system supporting human cognition and episodic memory^15^. Efficient suppression of DMN activity during frontoparietal control network engagement predicts superior attention performance^17^, while coordinated interactions among hippocampal and distributed cortical DMN nodes are associated with the subjective experience of remembering^15^.

Longitudinal human studies indicate that altered DMN functional connectivity predicts memory decline in both AD patients and cognitively healthy adults and these alterations are spatially correlated with progression of amyloid pathology, underscoring its role as an early marker of cognitive deterioration^18–21^. However, translational validation of network-level findings in preclinical models is limited^22–27^, and the cellular mechanisms underlying amyloid-associated network dysfunction are poorly understood.

Myelin, produced by oligodendrocytes, is essential for the rapid signal propagation, metabolic support and the temporal precision required for coordinated neural activity^28–30^. Oligodendrocyte precursor cells (OPCs), defined by expression of the receptor tyrosine kinase PDGFRα, maintain myelin integrity throughout life and serve as the primary source of remyelinating cells in the adult brain^31^. Beyond their role in myelin repair, neurons form synaptic contacts onto OPCs, which act as sensors of neuronal activity to regulate circuit development and remodeling^32,33^.

Late-myelinating brain regions are among the first to degenerate during normal aging^34–36^. Notably, the DMN regions most vulnerable to functional connectivity disruption in AD including the cingulate cortex and hippocampus overlap with these regions of aging-dependent preferential myelin loss ^37^, suggesting a mechanism linking myelin vulnerability to network dysfunction.

Given myelin’s fundamental role in regulating the conduction velocity and temporal synchrony of long-range axonal projections^31,38^, region-specific demyelination within DMN nodes could represent a direct cellular substrate of the functional connectivity alterations observed in AD.

Amyloid pathology further disrupts this system at multiple levels. Aβ accumulation exacerbates age-related myelin degeneration, impairing white matter integrity and reducing axonal conduction velocity and temporal synchrony. This disruption of network coherence contributes to neurodegeneration and cognitive decline^39–41^. Furthermore, amyloid pathology induces OPC dysfunction, driving OPC’s premature senescence, characterized by upregulation of cell cycle inhibitors (e.g., p16^INK4a^, p21^CIP1^), secretion of pro-inflammatory cytokines (the senescence-associated secretory phenotype, SASP), and reduced proliferative capacity^42^. OPCs associated with Aβ plaques exhibit aging-like phenotypes and contribute to neuroinflammatory cascades within the plaque microenvironment. Noteworthy, senolytic clearance of these cells reduces microglial reactivity and Aβ burden^43^. Conversely, oligodendrocytes and OPCs express amyloid precursor protein (APP) and the key components of the amyloidogenic pathway including BACE1, PSEN1, and PSEN2 at levels comparable to neurons and contribute to plaque development^44,45^. APP also plays a bidirectional role in myelin biology: amyloidogenic cleavage of APP by *β*-secretase and *γ*-secretase generates Aβ species that impair oligodendrocyte survival and disrupt myelin sheath formation, whereas non-amyloidogenic cleavage by α-secretase promotes OPC differentiation and myelin production^46^. In turn, myelin injury upregulates amyloidogenic APP processing, suggesting a self-amplifying cycle^47^. Together, accelerated myelin degeneration, impaired remyelination, and OPC senescence are poised to disrupt the structural basis of long-range neural synchrony. We hypothesize that region-specific alterations in myelin integrity and OPC function within DMN nodes constitute a key cellular mechanism linking amyloid pathology to large-scale network dysfunction.

Here, we address this gap by integrating ultra-high-field resting-state fMRI with quantitative triple immunofluorescence of MBP, PDGFRα, and Aβ plaques within the same DMN subregions across three disease stages in a humanized APP knock-in mouse model of AD^48^. This model recapitulates progressive Aβ plaque deposition and associated glial pathologies without the confound of APP overexpression. By combining functional network measures with spatially resolved cellular markers of myelin integrity and OPC density within a single experimental framework, we characterize the co-evolution of myelin architecture, OPC distribution, and large-scale network function at subregional resolution across AD progression. We demonstrate a biphasic trajectory of DMN dysfunction, consistent with observations in humans^49^, transitioning from early hypersynchrony to late-stage hyposynchrony. This pattern is accompanied by arrested myelin maturation and late-stage loss of myelin and OPCs, most pronounced in the anterior cingulate cortex (ACC) and hippocampal subfield CA3. MBP coverage positively predicted DMN amplitude in region-specific manner, whereas this coupling is selectively disrupted in the presence of amyloid pathology. Together, these data establish oligodendroglial dysfunction and myelin loss as key contributors to DMN failure in amyloidopathy.

## Results

### Biphasic trajectory of DMN in App^NL-G-F/NL-G-F^ mice across age groups

We examined DMN synchrony in APP knock-in mice across three different ages during the progression of amyloid pathology to characterize the evolution of large-scale network alterations. Network-level and ROI-to-ROI analyses revealed pronounced age-dependent reorganization of DMN synchrony. App^NL-G-F/NL-G-F^ mice exhibited increased DMN synchrony relative to App^NL/NL^ controls, most prominently within the ACC, S1, and CA1, revealing a phase of network *hypersynchrony* at the 3-months of age (Fig. 1A). By 7 months, this pattern shifted toward *hyposynchrony*, particularly in the ACC bilaterally and CA1 unilaterally, while S1 remained hypersynchronous, showing region-specific differences in the progression of DMN dysfunction relative to App^NL/NL^ during a period of advancing plaque burden (Supplementary Fig. 1). In aged mice with saturated plaque burden (+1 year), genotype-dependent differences in DMN synchrony were markedly attenuated, with loss of hypersynchrony and only sparse residual hyposynchrony persisting in the ACC of App^NL-G-F/NL-G-F^ mice compared to App^NL/NL^.

**Figure 1.**
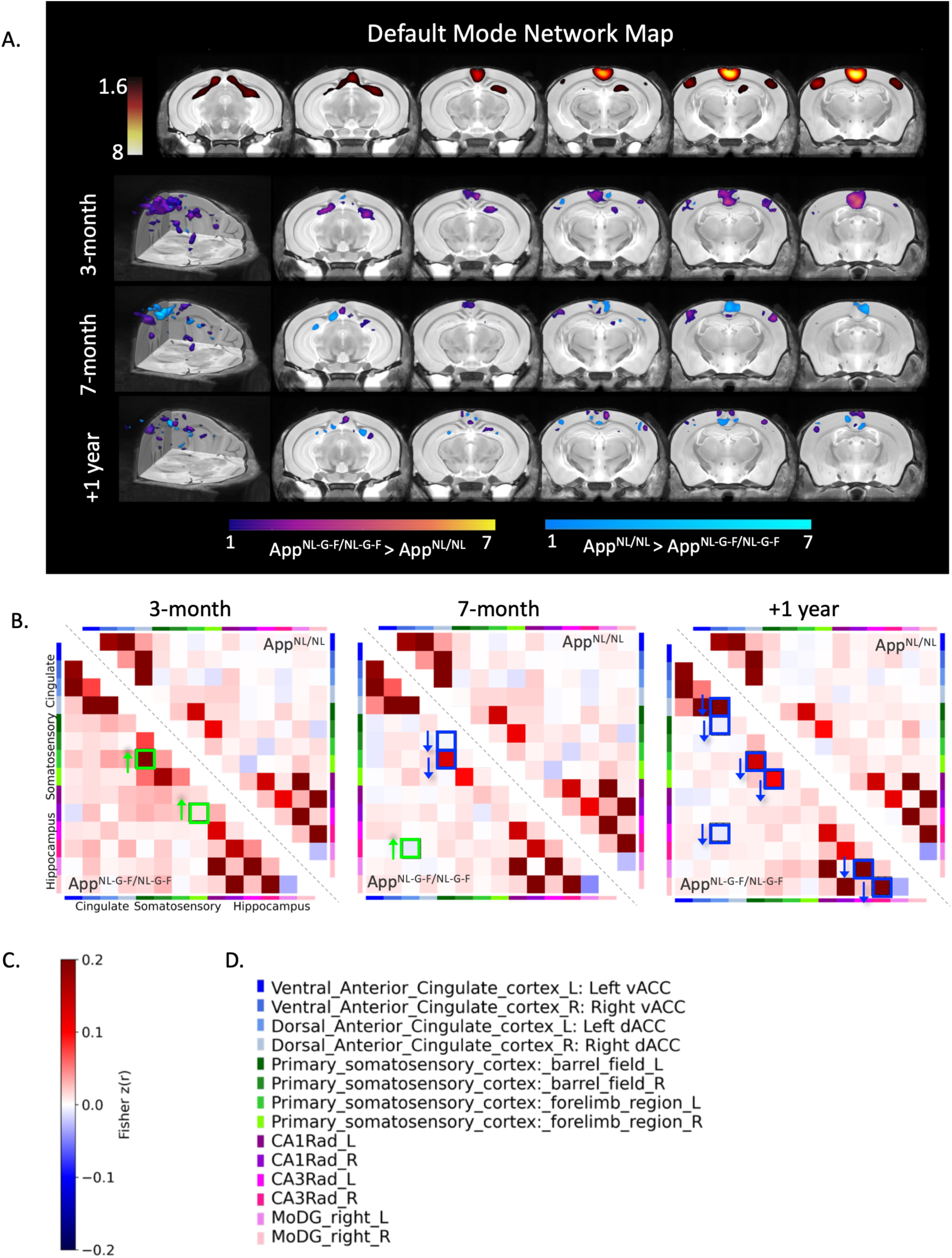
Default Mode Network functional connectivity in APP knock-in mice across ages. **(A)** Group-level DMN spatial map derived from independent component analysis (ICA) of rs-fMRI data (top) and voxelwise statistical comparisons of DMN connectivity between App^NL-G-F/NL-G-F^ and App^NL/NL^ mice at 3 months, 7 months, and +1 year. Purple–yellow scale indicates regions where App^NL-G-F/NL-G-F^ > App^NL/NL^, whereas cyan–blue scale indicates regions where App^NL/NL^ > App^NL-G-F/NL-G-F^. **(B)** Functional connectivity matrices depicting pairwise correlations between DMN subregions at 3 months, 7 months, and +1 year of age. Green-outlined boxes denote significant between-genotype differences (App^NL-G-F/NL-G-F^, bottom triangle vs. App^NL/NL^, upper triangle; split across the diagonal); blue-outlined boxes with arrows denote significant within-genotype age-dependent changes. For between-genotype comparisons (green boxes), upward arrows (↑) indicate increased connectivity in App^NL-G-F/NL-G-F^ relative to App^NL/NL^. For within-genotype comparisons (blue boxes) downward arrows (↓) indicate decreased connectivity relative to the preceding timepoints in App^NL-G-F/NL-G-F^ group. **(C)** Color scale for connectivity matrices, representing Fisher z-transformed correlation coefficients (r), ranging from negative (blue) to positive (red) connectivity. **(D)** Color-coded region legend identifying the DMN subregions represented in the connectivity matrices, corresponding to regions also quantified by immunofluorescence.

While ICA revealed the spatial distribution of synchrony changes, ROI-to-ROI functional connectivity analysis within the DMN allowed quantification of the changes (Fig. 1B). At 3-months, App^NL-G-F/NL-G-F^ mice exhibited increased connectivity with intra-cortico-cortical and cortical-hippocampal connections including CA1–S1 and S1(barrel field)–S1(forelimb region) relative to App^NL/NL^ controls. At 7-month, App^NL-G-F/NL-G-F^ mice exhibited a mixed pattern of functional connectivity changes along with advanced amyloid pathology (Supplementary Fig1.). Particularly, inter-frontal-hippocampal (Right CA3 and Left Dorsal ACC) synchrony was increased in App^NL-G-F/NL-G-F^ mice compared to App^NL/NL^ controls, while inter- and intra-cortical synchrony involving S1 was decreased relative to the same App^NL-G-F/NL-G-F^ mice at 3 months of age. At +1 year, within-genotype reductions in functional connectivity had further progressed in App^NL-G-F/NL-G-F^, affecting several DMN nodes. These included intra-frontal-cortical (Left dorsal ACC-S1), intra-hippocampal (Left CA3-DG), and intra frontal-hippocampal pathways (Left CA3–ACC), along with inter-frontal changes in the dorsal ACC (*p* < 0.05). In contrast, App^NL/NL^ control mice exhibited no significant changes in functional connectivity between age groups.

### Progressive demyelination and region-dependent OPC loss in App^NL-G-F/NL-G-F^ mice

The availability of OPCs is a key determinant of white matter integrity, neural circuit function and repair capacity. To link these cellular properties to DMN functional connectivity, we quantified MBP, a marker of mature myelin and myelinating oligodendrocytes, and PDGFRα, a marker of oligodendrocyte precursor cells (OPCs) (Figs. 2–3), alongside Aβ plaques (Supplementary Fig. 1) within the same DMN subregions examined in the functional connectivity analysis (Fig. 1) across all three age groups. For the 7-month and +1 year cohorts, brain tissue was collected from the same animals used for MRI, enabling direct spatial and within-subject correspondence between myelin-related structural changes and DMN functional connectivity during plaque progression.

**Figure 2.**
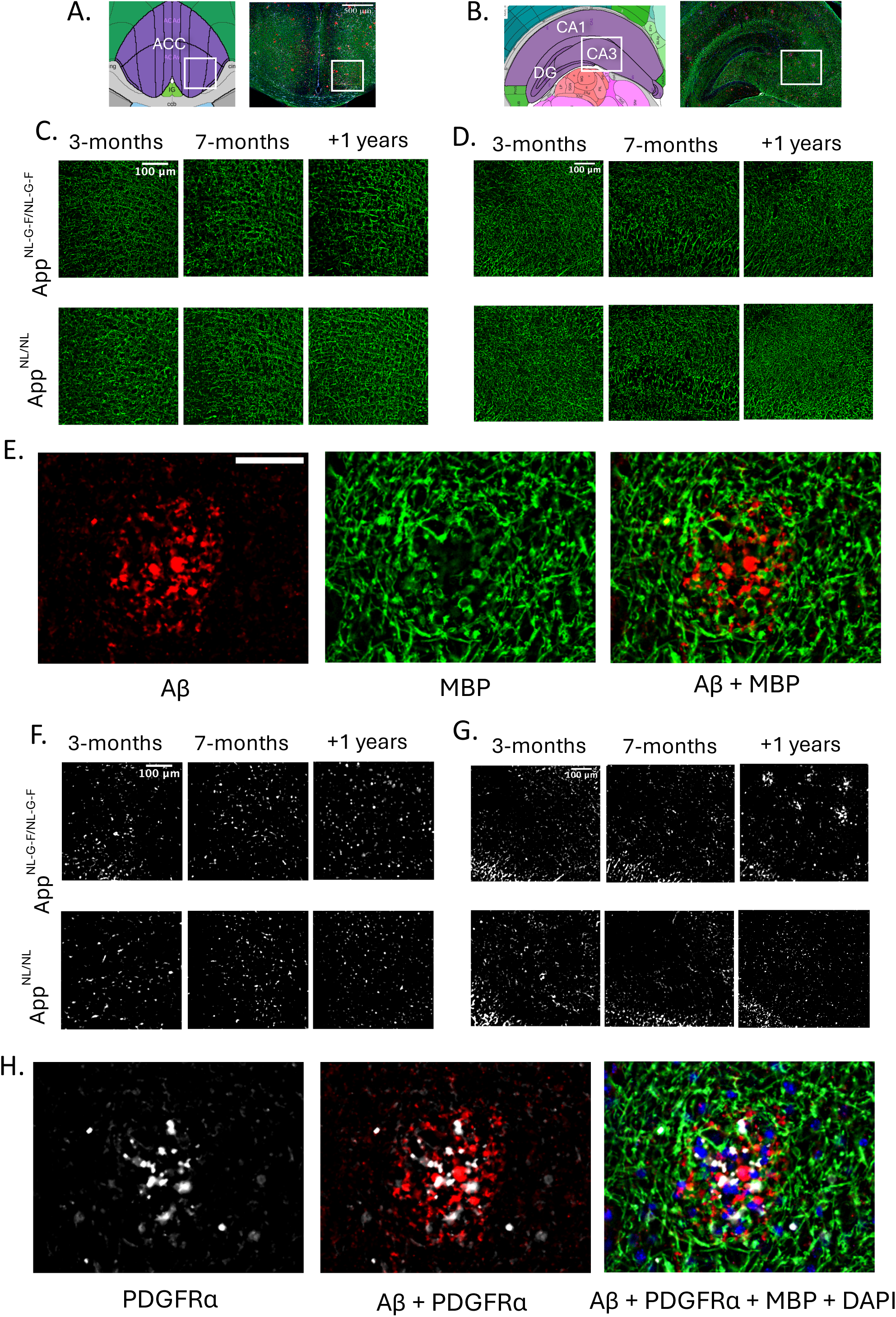
MBP and PDGFRα expression across cortical and hippocampal regions in APP knock-in mice across age groups. **(A–B)** Representative brain parcellations from the Allen Brain Atlas^50^ and fluorescence images showing the cortical (A) and hippocampal (B) regions of interest. White boxes indicate areas displayed in C–D and F–G. **(C–D, F–G)** Representative MBP (green) (C-D) and PDGFRα (white) (F-G) immunostaining in the cortex (C, F) and hippocampus (D, G) of App^NL-G-F/NL-G-F^ (top) and App^NL/NL^ (bottom) mice at 3 months (3m), 7 months (7m), and +1 year (+1yr). Scale bar = 100 µm. **(E)** Representative close-up images from 7 months old, male App^NL-G-F/NL-G-F^ mouse in CA3 showing Aβ (red), MBP (green), and merged overlay, illustrating absence of MBP expression in the vicinity of Aβ plaques. Scale bar = 40 µm. **(H)** Fluorescence images showing PDGFRα (white/gray), Aβ (red), MBP (green), and nuclei (DAPI; blue), illustrating OPC enrichment within the periplaque zone in areas of myelin loss.

**Figure 3.**
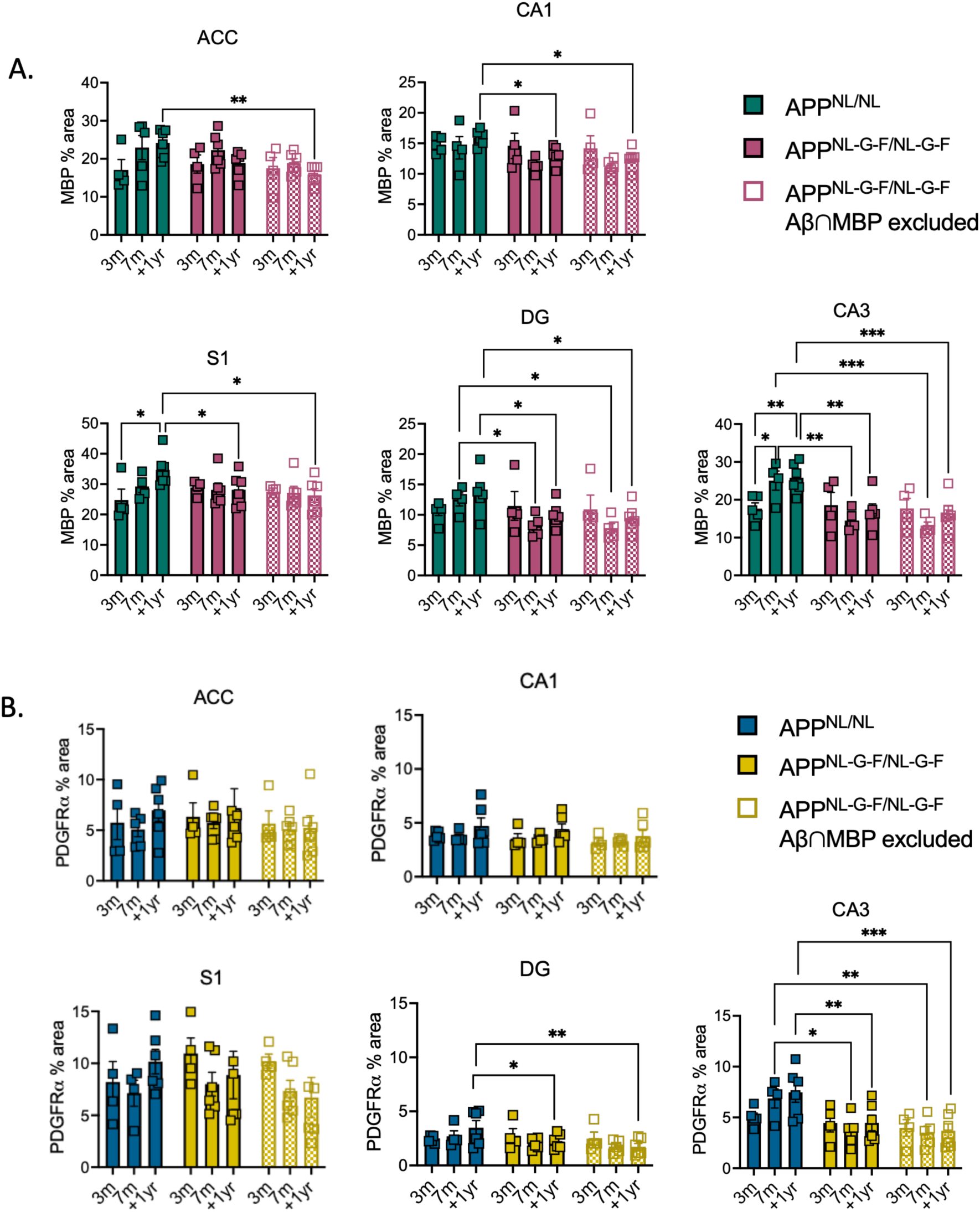
Quantification of MBP and PDGFRα coverage across cortical and hippocampal regions in APP knock-in mice across age groups. **(A)** MBP immunofluorescence signal (% area) across ACC, CA1, CA3, S1, and DG at 3 months (3m), 7 months (7m), and +1 year (+1yr) in App^NL/NL^ (teal), App^NL-G-F/NL-G-F^ (pink), and App^NL-G-F/NL-G-F^ excluding areas where Aβ-colocalized with MBP (App^NL-G-F/NL-G-F^ w/o Aβ, hatched pink). **(B)** PDGFRα immunofluorescence signal (% area) across the same regions and timepoints in App^NL/NL^ (blue), App^NL-G-F/NL-G-F^ (yellow), and App^NL-G-F/NL-G-F^ excluding areas where Aβ colocalized with PDGFRα (hatched yellow). Individual data points shown as squares. Data were analyzed by two-way ANOVA with Tukey’s correction for multiple comparisons. \**p* < 0.05, \*\**p* < 0.01, \*\*\**p<0.001*.

To characterize the association between oligodendrocyte lineage markers and Aβ plaques, we quantified the colocalization of MBP and PDGFRα with Aβ deposits (Supplementary Fig. 2) and calculated MBP and PDGFRα signal area percentages both with and without the colocalized fraction (Fig. 3). This approach is justified on two grounds: first, App^NL/NL^ control mice are devoid of Aβ plaques and therefore represent a plaque-independent baseline; and second, it remains unclear whether areas with MBP and PDGFRα signal colocalized with Aβ deposits retain normal functional properties.

First, we calculated total MBP area fraction across ROIs, including areas colocalized with Aβ plaques, and compared App^NL-G-F/NL-G-F^ and App^NL/NL^ mice using a two-way ANOVA (genotype × timepoint) followed by Tukey’s post-hoc test. Total MBP coverage was reduced in App^NL-G-F/NL-G-F^ mice relative to App^NL/NL^ controls in S1 (+1yr: *p* = 0.0420), CA1 (+1yr: *p* = 0.0397), DG (7m: *p* = 0.0275; +1yr: *p* = 0.0321), and CA3 (7m: *p* = 0.0016; +1yr: *p* = 0.0022), whereas ACC did not reach statistical significance (+1yr: *p* = 0.0690). Notably, CA3 and DG showed MBP loss as early as 7 months, indicating early regional vulnerability.

We subsequently computed MBP coverage after excluding colocalized areas in App^NL-G-F/NL-G-F^ mice and compared to App^NL/NL^ mice, with a separate two-way ANOVA (genotype × timepoint) with Tukey’s post-hoc test. This analysis revealed significant MBP coverage area loss across all ROIs at +1 year (ACC: *p* = 0.0042; S1: *p* = 0.0102; CA1: *p* = 0.0170; CA3: *p* = 0.0007; DG: *p* = 0.0156), with CA3 and DG also showing reductions at 7 months (CA3: *p* = 0.0003; DG: *p* = 0.0184), revealing progressive demyelination in App^NL-G-F/NL-G-F^ mice. Notably, there was no reduction in total MBP area in ACC when all colocalized areas were included; the signal area loss emerged once Aβ-colocalized MBP was accounted for, implicating Aβ association as a key contributor to myelin loss in this region.

Notably, App^NL/NL^ mice exhibited significant age-dependent *increases* in MBP coverage in S1, and CA3 between 3 months and +1 year (S1: *p* = 0.0193; CA3: *p* = 0.0095), and between 3 and 7 months in CA3 (*p* = 0.0358), demonstrating that myelination in these regions continues to mature during normal aging in the absence of amyloid plaque pathology. We do not observe this continuing maturation in App^NL-G-F/NL-G-F^ mice, highlighting the extent to which Aβ plaque burden disrupts normal age-dependent myelin maturation in these regions.

App^NL-G-F/NL-G-F^ mice showed significantly reduced PDGFRα coverage compared to App^NL/NL^ mice at both 7 months and +1 year in CA3, regardless of whether Aβ-colocalized areas were included (total: 7m *p* = 0.0156, +1yr *p* = 0.0086; excluding colocalization: 7m *p* = 0.0083, +1yr *p* = 0.001), indicating a progressive reduction in OPC density. A significant difference was also observed in DG at +1 year when colocalized areas were included (*p* = 0.0419) and excluded (*p* = 0.0066), consistent with the effect observed for MBP. No significant changes in PDGFRα coverage were detected between genotypes or across age in ACC, S1, or CA1.

**Figure 3.**
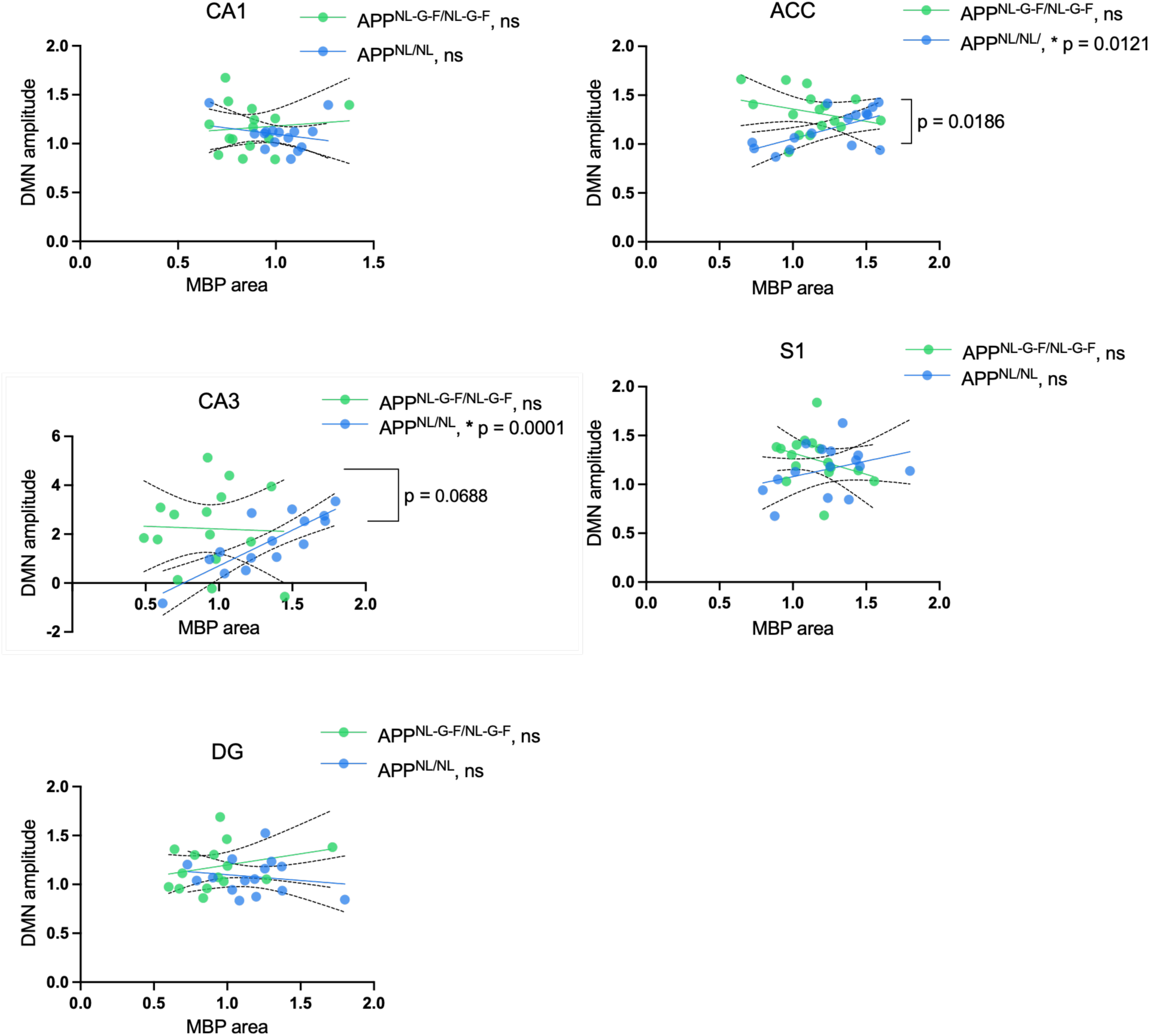
MBP-DMN coupling is disrupted in APP knock in mice in a region-dependent manner. Scatter plots showing the relationship between normalized MBP coverage (x-axis; MBP % area normalized to 3-month App^NL/NL^) and normalized DMN amplitude (y-axis). Lines represent simple linear regression fits with 95% confidence intervals (dashed). For App^NL-G-F/NL-G-F^ mice, MBP values excluding Aβ-colocalized areas were used. Green = App^NL-G-F/NL-G-F^; blue = App^NL/NL^.

### MBP-DMN coupling breakdown in App^NL-G-F/NL-G-F^ mice in a region-dependent manner

Lastly, we examined whether myelin integrity is associated with DMN strength in the subregions (ROIs) comprising the DMN. The dual regression maps were used to assess how strongly the BOLD signal in each ROI fluctuates in synchrony with the DMN time course, hereafter referred to as DMN amplitude.

In App^NL/NL^ mice, MBP coverage was positively associated with DMN amplitude in both ACC (*p* = 0.0121, slope = 0.400, R² = 0.395) and CA3 (*p* = 0.0001, slope = 2.914, R² = 0.692), demonstrating that in the absence of amyloid pathology, greater myelin integrity is associated with higher amplitude within DMN in these regions (Fig. 3). No association between MBP coverage and DMN amplitude was detected in App^NL-G-F/NL-G-F^ mice in any region (ACC: slope = −0.254, *p* = 0.288, R² = 0.080; CA3: slope = −0.220, *p* = 0.898, R² = 0.001), indicating that this relationship is disrupted in the presence of amyloid plaque pathology. Comparison of regression lines between genotypes revealed a difference in slopes in ACC (F = 6.278, *p* = 0.0186), reflecting a positive MBP-DMN relationship in App^NL/NL^ mice that is reversed in App^NL-G-F/NL-G-F^ mice. No associations or between-genotype differences were detected in S1, DG, or CA1, demonstrating that MBP-DMN coupling is region-specific. These findings demonstrate that myelin integrity is coupled to DMN amplitude in control mice in ACC and CA3, and that this coupling is disrupted in the presence of amyloid pathology, implicating demyelination as a potential contributor to structure-function coupling disruption in App^NL-G-F/NL-G-F^ mice.

## Discussion

The evolution of functional network alterations during amyloid plaque progression is well established^51–56^ and could serve as an alternative biomarker of AD progression, pending a better understanding of the underlying biology. By integrating ultra-high field rs-fMRI with triple immunofluorescence microscopy, we investigated the contribution of OPCs and MBP to network alterations under normal conditions and across amyloid pathology progression. We observe that advancing amyloid pathology drives a biphasic reorganization of DMN functional connectivity, from early hypersynchrony to hyposynchrony, concurrent with disrupted myelin maturation, progressive demyelination and OPC loss. We found that myelin coverage, which is positively coupled to DMN strength in control mice, becomes decoupled in the presence of amyloid pathology in a region-specific manner. This establishes amyloid-driven, region-specific myelin dysfunction and demyelination as a cellular substrate for DMN failure in a humanized APP knock-in mouse model of Alzheimer’s disease.

The hypersynchrony observed in App^NL-G-F/NL-G-F^ mice with amyloidopathy compared to age-matched App^NL/NL^ control mice at 3 months coinciding with initial Aβ plaque deposition is consistent with reports of network hyperconnectivity in preclinical AD in both humans and transgenic mouse models^51–56^. Within a network control framework, reserve can be conceptualized as the brain’s residual capacity to reconfigure functional connectivity in order to sustain cognitive performance despite accumulating pathology, a capacity thought to be instantiated through compensatory hyperconnectivity in early disease stages (Medaglia, Pasqualetti, Hamilton, Thompson-Schill, & Bassett, 2017; Yoo et al., 2015). However, compensatory hyperconnectivity could further exacerbate the pathology. Recent evidence indicates that Aβ-induced hyperconnectivity can causally accelerate tau spread by increasing functional connectivity between tau epicenters and tau-vulnerable posterior regions, thereby mediating faster tau accumulation (Roemer-Cassiano et al., 2025). This observation is compatible with a model in which early hypersynchrony represents a dual-edged network state, one that may transiently support cognitive maintenance while inadvertently also facilitating the activity-dependent propagation of pathology. At this stage, App^NL-G-F/NL-G-F^ mice perform comparably to controls on conventional spatial memory paradigms including Morris water maze^57^ and Barnes maze tasks^58^ at 3-4 months old, consistent with a presymptomatic stage of AD. However, at 4-5 months old, these mice do show impairment in touchscreen-based pattern separation and paired associative memory (but not hippocampus-independent touchscreen tasks like visual discrimination)^59^. The sensitivity of the touchscreen-based pattern separation and paired associative memory tasks to early AD-related pathology has been attributed to their dependence on hippocampal subregional circuits, particularly the entorhinal cortex–DG–CA3– CA1 network which represent primary sites of Aβ pathology at this disease stage^59^. DMN hypersynchrony may be insufficient to disrupt performance on tasks amenable to distributed cortical networks, but sufficient to impair the subcortical hippocampal-specific computations that pattern separation and associative memory demand^59^. This suggests that combining rs-fMRI network analysis with touchscreen-based hippocampal-specific cognitive testing may provide a sensitive early window for AD detection.

DMN synchrony shifted toward hyposynchrony in ACC and CA1 while S1 remained hypersynchronous in App^NL-G-F/NL-G-F^ mice by 7-months, revealing region-specific differences in the pace of network dysfunction during a period of advancing plaque burden. The App^NL-G-F/NL-G-F^ considered as one of the aggressive amyloid pathology models of AD that carry the Arctic, Swedish and Beyreuther/Iberian mutations. Cortical Aβ plaque deposition begins as early as 2 months of age^48^, saturates at 7-months old (Supplementary Fig.1C) and neuroinflammatory responses are observed including microgliosis and astrocytosis at 9-month^48^. Emergence of the cognitive deficits in conventional behavioural paradigms is also observed at 7-month old App^NL-G-F/NL-G-^ mice, including impairments in fear learning^60^, spatial learning^58^ (while retained memory) as well as touchscreen-based visual discrimination^61^, which were reported as intact in earlier timepoints. This suggests that the onset of cognitive impairment on several domains is not contemporaneous with the onset of amyloid pathology but rather tracks the collapse of the compensatory network state and advanced amyloid burden. Therefore, the transition from hyper to hyposynchrony may constitute a functionally critical tipping point where amyloid toxicity reaches critical levels, leading to a situation where network reserve that underlies plasticity is exhausted, resulting in broader cognitive deterioration beyond hippocampus sensitive domains. This hypothesis has direct implications for the timing of diagnostic features (e.g. conversion from mild cognitive impairment to AD).

Large-scale integrative brain networks such as the DMN are critically important for cognition and fundamentally dependent on the structural integrity of the axons that connect their constituent nodes. Myelin is the primary determinant of axonal conduction velocity, temporal synchrony, and the metabolic support of long-range projections^38^. Beyond its structural role, myelination is a dynamic, experience-dependent process that continues well into late adulthood, with OPCs responding to neuronal activity to generate new myelin and refine circuit function across the lifespan^13,31,62–68^. Consistent with this view, our data reveal that APPKI1 mice exhibit significant age-dependent increases in MBP coverage in ACC, S1, and CA3 between 3 months and +1 year, reflecting the normal continuation of myelin maturation in these regions in the absence of amyloid pathology. This process is entirely absent in App^NL-G-F/NL-G-F^ mice, indicating that amyloid pathology actively arrests the normal trajectory of age-dependent myelin maturation, producing a progressive, region-dependent failure of myelin maturation that emerges with plaque burden.

In addition to the failure of myelin maturation, the progressive co-evolution of demyelination and alterations in DMN synchrony across the pathological stages raises the question of whether these processes are mechanistically linked or represent parallel but independent consequences of amyloid pathology. Our linear regression analysis directly addresses this question by demonstrating that MBP coverage is positively associated with DMN amplitude in control mice in both ACC and CA3, establishing myelin coverage as a structural predictor of DMN strength in the healthy brain, and the absence of this coupling in App^NL-G-F/NL-G-F^ mice provides evidence that demyelination disrupts the structural basis of functional network dynamics in the context of amyloid pathology. We hypothesize that the recruitment of existing MBP and OPCs to sites of amyloid injury redirects their structural and signaling contributions away from the maintenance of network synchrony, severing the normal correspondence between myelin distribution and functional network strength^69–71726173,74^. Together, these findings position the oligodendroglial lineage as a critical determinant of functional network integrity in AD and establishes a clear spatiotemporal framework in which early DMN hypersynchrony precedes detectable myelin loss and cognitive symptoms in AD, followed by progressive demyelination and OPC depletion in hippocampal subfields that converges with late-stage network hyposynchrony and myelin-network decoupling. Future studies leveraging rs-fMRI in awake mouse models may directly test whether targeted oligodendroglial therapeutics alone or in combination with amyloid-targeting drugs can rescue or reverse network dysfunction and cognitive decline, providing causal evidence that functional network alterations are both a readout of and a modifiable contributor to AD-related pathology.

## Methods

### Animals

The App^NL-G-F/NL-G-F^ knock-in mouse model of AD, previously introduced by Saito et al. (2014), was used as the amyloid-bearing line^48^. This model harbours three familial AD mutations in the endogenous App gene: the Swedish (KM670/671NL), Arctic (E693G), and Beyreuther/Iberian (I716F) mutations, which together increase total Aβ production, promote Aβ oligomerization, and elevate the Aβ42/Aβ40 ratio, respectively. The App^NL/NL^ knock-in line, which carries only the humanized Swedish mutation was used as controls. This line mirrors the APP processing alterations of App-^NL-G-F/NL-G-F^ but do not develop amyloid plaques and isolates the effects of plaque deposition from those of altered Aβ production. All animals were obtained from the Prado Laboratory at Western University. All animal procedures were performed in accordance with the Canadian Council on Animal Care guidelines and approved Animal Use Protocols at the University of Western Ontario (Protocol # 2020-162). All mice had unrestricted access to food and water.

### Experimental Design

Three age groups were examined: 3 months (3m), 7 months (7m), and +1 year (+1yr). +1yr groups consisted of animals aged between 13.8-18.1 months. A longitudinal design was employed for the 3m–7m cohort, in which the same animals underwent MRI at both timepoints (App^NL-G-F/NL-G-F^: n = 9; App^NL/NL^: n = 6), with the 7m animals subsequently used for immunofluorescence (IF) analysis (n = 4–6 per genotype). A separate cohort was used for 3m IF studies (n = 4–5 per genotype). For the +1yr cohort, MRI (App^NL-G-F/NL-G-F^: n = 11; App^NL/NL^: n = 9) and IF (n = 6 per genotype) were performed on the same animals.

### MRI Acquisition

All MRI procedures were conducted on a 9.4T MRI magnet (Varian NMR, USA) interfaced to a AV3HD or Neo MRI console running Paravision 360 (Bruker, Germany) located in the Centre for Functional and Metabolic Mapping at Western University. All animals were anesthetized with 1.5-2% isoflurane during the scans.

#### T2-TurboRARE

The T2-weighted anatomical images were acquired using a TurboRARE2D pulse sequence (16 averages, 31 slices, slice thickness = 0.5 mm, FOV 19.2 × 15 mm^2^, matrix size 128×100, in-plane resolution = 0.15 × 0.15 mm^2^, TE = 40 ms, TR = 7000 s, echo spacing = 10 ms, and RARE factor 8).

#### Rs-fMRI

Resting state fMRI (600 volumes x 5 runs) were acquired using a 2-dimensional echo-planar imaging sequence (EPI2D) with parameters: TR = 1500 ms, TE = 12 ms, and flip angle = 60°. Each functional volume comprised of 31 slices with an in-plane resolution of 0.3 × 0.3 mm^2^ and a slice thickness of 0.5 mm. The field of view was 19.2 x 9.6 mm^2^, matrix size was 64 x 32, and bandwidth = 250kHz.A single loop RF coil was used for the scans.

### MRI Analysis

#### Preprocessing

All data preprocessing and registration steps performed using RABIES (Rodent Automated BOLD Improvement of EPI Sequences) version 0.4.8 (https://rabies.readthedocs.io/en/stable/)^69^. The pipeline included robust correction procedures for both structural and functional inhomogeneities to reduce biases related to signal distortions and intensity inconsistencies arising from magnetic field inhomogeneity, coil sensitivity variations, and other acquisition-related factors.

To isolate the BOLD signal from non-neuronal sources, nuisance regression was conducted to remove signal contributions from white matter, cerebrospinal fluid (CSF), major vascular structures and six motion parameters. ICA-AROMA was applied to remove further effects of motion. A high-pass temporal filter with a cutoff frequency of 0.01 Hz was applied to eliminate low-frequency signal drifts.

Following denoising, each BOLD scan was registered to its corresponding anatomical scan. The high-resolution anatomical images were subsequently aligned to the DSURQE standard brain atlas^70^ and the resulting transformation matrix was also applied to the corresponding BOLD data to enable consistent brain parcellation within the RABIES pipeline. This atlas-based parcellation allowed for the extraction of regional timeseries and the investigation of whole-brain functional connectivity across animals.

#### Network-level and ROI-ROI analysis for the Default mode network

To investigate connectivity between the nodes constituting the DMN, we used independent component analysis (ICA) implemented in the RABIES pipeline. Group-level ICA was performed to derive data-driven spatial components from resting-state BOLD time series. Individualized connectivity estimates were then obtained via dual regression, which models scan-specific versions of each group-level component through two sequential linear regressions. In the first step, component time courses were estimated for each subject using multivariate linear regression; in the second step, subject-specific spatial maps were reconstructed from these time courses. Time courses from the first regression step were variance-normalized using the root mean square of each component time course prior to the second regression step to allow unbiased estimation of connectivity amplitudes; then in the second step, the normalized time courses were regressed against each subject’s BOLD timeseries to derive subject-specific spatial maps, in which each voxel’s value reflects the strength of its coupling to the DMN in that subject. The resulting DMN component maps captured subject-level variation in both network amplitude and spatial configuration, enabling statistical comparisons across genotypes and age groups.

Statistical differences in dual regression maps between genotypes across age groups within the DMN (Fig. 1A) were assessed using a permutation-based GLM (5,000 permutations) implemented in PALM/FSL (alpha119; MATLAB R2023b), producing z-statistic outputs with voxelwise false discover rate (FDR) control and contrast-wise correction (-zstat -n 5000 -fdr - corrcon)^71^.

To quantify how network-level alterations were reflected in interregional communication, ROI-based functional connectivity matrices were derived from subject-level Pearson correlation coefficients (r) computed from cleaned time series across DMN subregions, including cingulate (ventral and dorsal), hippocampal (CA1, CA3, and DG), and primary somatosensory (barrel field and forelimb) nodes (S1) (Fig. 1B). ROI selection was constrained to regions also sampled in the immunofluorescence experiments, enabling direct spatial correspondence between functional connectivity measures and cellular and molecular analyses in those regions. Correlation coefficients were subsequently Fisher r-to-z transformed to improve normality prior to group-level comparisons. All statistical inference for pairwise differences was performed using a permutation-based framework implemented in PALM/FSL and corrected for multiple comparisons using FDR^71^.

### Immunofluorescence

#### Brain tissue preparation

Mice were anesthetized with ketamine (1-1.5mg/kg) and perfused with 4% cold paraformaldehyde (PFA) transcardially after an initial flush with phosphate-buffered saline (PBS). Brains were post-fixed in 4% PFA at 4°C for 24 hours, embedded in optimal cutting temperature (OCT) compound, and cut into 40-μm-thick sections coronally using a cryostat microtome. The slices were kept in cryoprotectant at 4°C for storage. Free-floating sections were washed with phosphate buffered saline (PBS) (3×5minutes), blocked with PBS + Triton-X100 + 4% normal goat serum (NGS) for one hour at room temperature, then mounted on the slides, 4 sections per mice was used.

#### Immunostaining protocol and quantification

Slide-mounted brain sections were subjected to antigen retrieval in 10 mM sodium citrate buffer (pH 6.1) at 85 °C for 30 min, cooled on ice for 45 min, and rinsed in Tris-buffered saline (TBS). Sections were permeabilized with TBS containing 0.3% Triton X-100, followed by blocking in TBS supplemented with 2% bovine serum albumin (BSA), 2% normal goat serum, 5% donkey serum, and 0.3% Triton X-100 for 90 minutes at room temperature.

Primary antibodies were diluted in blocking buffer consisting of 2% bovine serum albumin (BSA), 2% normal goat serum, 5% donkey serum, and 0.3% Triton X-100 in tris-buffered saline (TBS), clarified by centrifugation for 5 min at 15,000 g, and applied overnight at 4°C. The following primary antibodies were used: rabbit anti-Aβ1–42 (1:300, Abcam Cat# ab201061) for amyloid-beta plaque, mouse anti-MBP (1:1000, Invitrogen, clone 7G7, Cat# MA5-47469) for myelin basic protein and rat anti-PDGFRα (1:500, Abcam ab90967, clone APA5) for OPC expression.

After primary incubation, sections were washed in TBS (2 × 5 min) and incubated for 2.5 h at 4 °C with species-specific secondary antibodies diluted in TBS containing 1% BSA, 2% normal goat serum, and 2.5% donkey serum. Secondary antibodies used were: donkey anti-rabbit Alexa Fluor 647 (1:500, Invitrogen, Cat# A31573), goat anti-mouse Alexa Fluor 488 (1:500, Invitrogen, Cat#A11001), and goat anti-rat Alexa Fluor 594 (1:500, Invitrogen, Cat# A11007). Nuclei were counterstained with DAPI. Sections were then coverslipped and the edges sealed with nail polish before storage at 4 °C in the dark.

Fluorescence imaging was performed using a Leica MICA microscope at 20× magnification with identical acquisition settings across all samples. Image processing and quantification were conducted in Fiji/ImageJ (NIH). Four sections per animal were analyzed and values averaged, with 4–6 mice per group. For the anterior cingulate cortex, primary somatosensory cortex, and hippocampal subregions, region-of-interest (ROI) boundaries were manually defined for each section. All images were Gaussian filtered, thresholded, and binarized, and individual channel masks were multiplied to generate intersection masks representing colocalization pairs. Colocalized and individual area fractions were subsequently quantified within the defined ROIs.

### MBP-DMN Coupling Analysis

To examine the coupling of myelin integrity and DMN connectivity, MBP immunofluorescence signal (% area) and DMN amplitude derived from dual regression were correlated across corresponding ROIs. For App^NL-G-F/NL-G-F^ mice, MBP values excluding the Aβ-colocalized areas were used to isolate myelin signal that was independent of plaque-associated MBP. Prior to analysis, both MBP % area and DMN amplitude values were normalized to the mean of the 3-month App^NL/NL^ group to account for inter-cohort variability and enable cross-timepoint comparisons on a common scale. Simple linear regression was performed for each ROI incorporating both genotypes simultaneously, with genotype included as a categorical variable to enable direct comparison of regression slopes and intercepts between App^NL/NL^ and App^NL-G-F/NL-G-F^ mice. All analyses were performed in GraphPad Prism.

## Acknowledgements

We thank all members of the Centre for Functional and Metabolic Mapping (CFMM), Western University, for their contributions to this work, A. Li and A. Eed for technical assistance with MRI experiments, and M. Bellyou for animal preparation prior to MRI scanning. We also thank J. Fan and C. Fodor for their assistance in gathering animals used in this study.

