## Supplementary Figures for "Myelin maturation failure and oligodendrocyte precursor loss underlie default mode network disruption in a humanized APP knock-in model of Alzheimer’s disease"

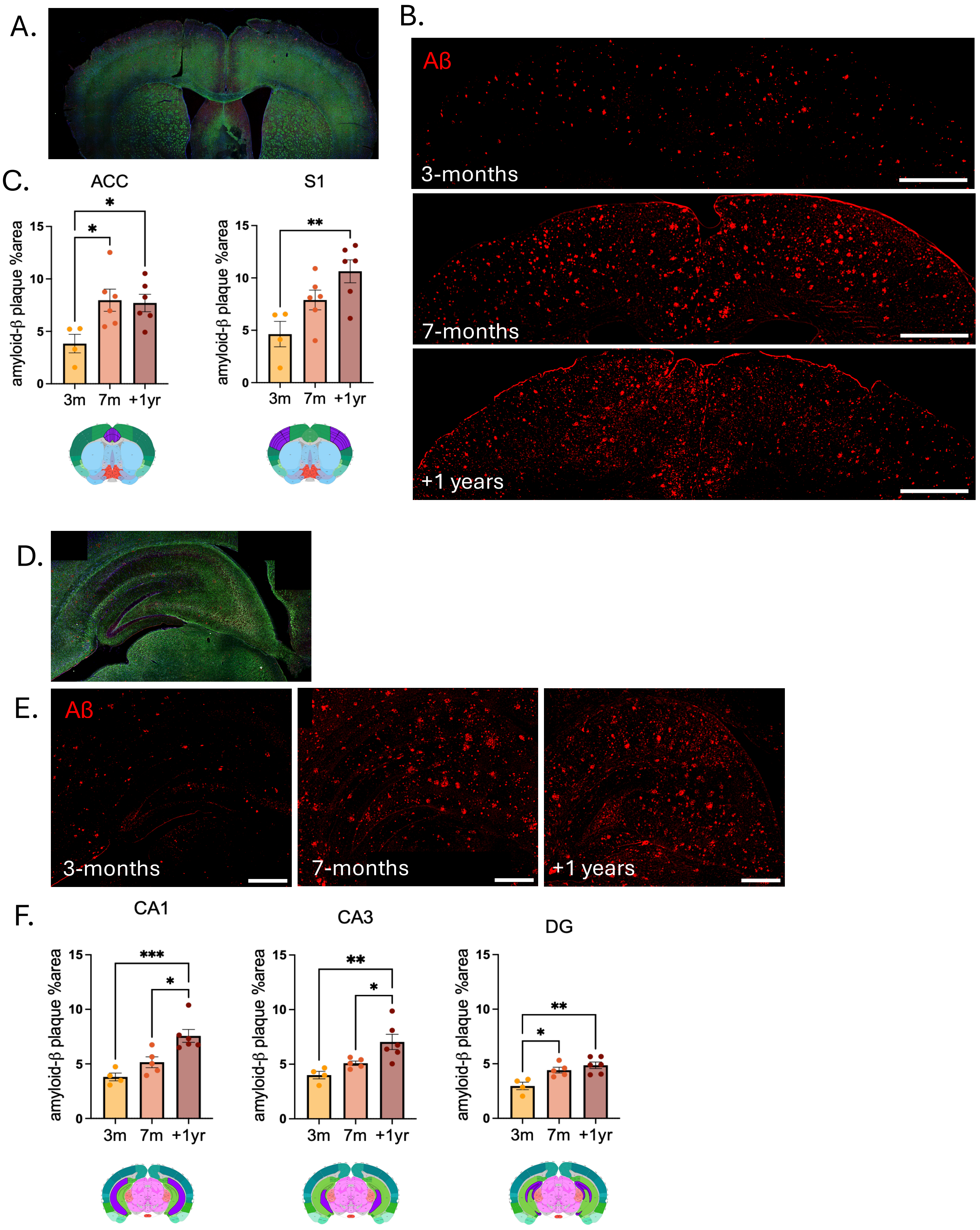
A.
B.
Aβ
3-months
7-months
+1 years
C.
ACC
S1
amyloid-β plaque %area
3m 7m +1yr
D.
E.
Aβ
3-months
7-months
+1 years
F.
CA1
CA3
DG
amyloid-β plaque %area
3m 7m +1yr

**Supplementary Figure 1. Progressive amyloid-β plaque deposition across cortical and hippocampal regions in APPKI3 mice**

(A) Representative merged fluorescence image of a coronal cortical section used per region. (B) Representative immunofluorescence images of Aβ plaque burden (red) in the cortex of $App^{NL\text{-}G\text{-}F/NL\text{-}G\text{-}F}$ mice at 3 months, 7 months, and ≥1 year. Scale bars = 500 µm. (C) Quantification of Aβ plaque % area in the ACC and S1 across age groups in $App^{NL\text{-}G\text{-}F/NL\text{-}G\text{-}F}$ mice. (D) Representative merged fluorescence image of a coronal hippocampal section. (E) Representative immunofluorescence images of Aβ plaque burden (red) in the hippocampus of $App^{NL\text{-}G\text{-}F/NL\text{-}G\text{-}F}$ mice at 3 months, 7 months, and +1 year. Scale bars = 500 µm. (F) Quantification of Aβ plaque % area in CA1, CA3, and DG across age groups. Data are presented as mean ± SEM. Statistical comparisons were performed using one-way ANOVA with Tukey's correction for multiple comparisons. *$p < 0.05$, **$p < 0.01$, ***$p < 0.001$.

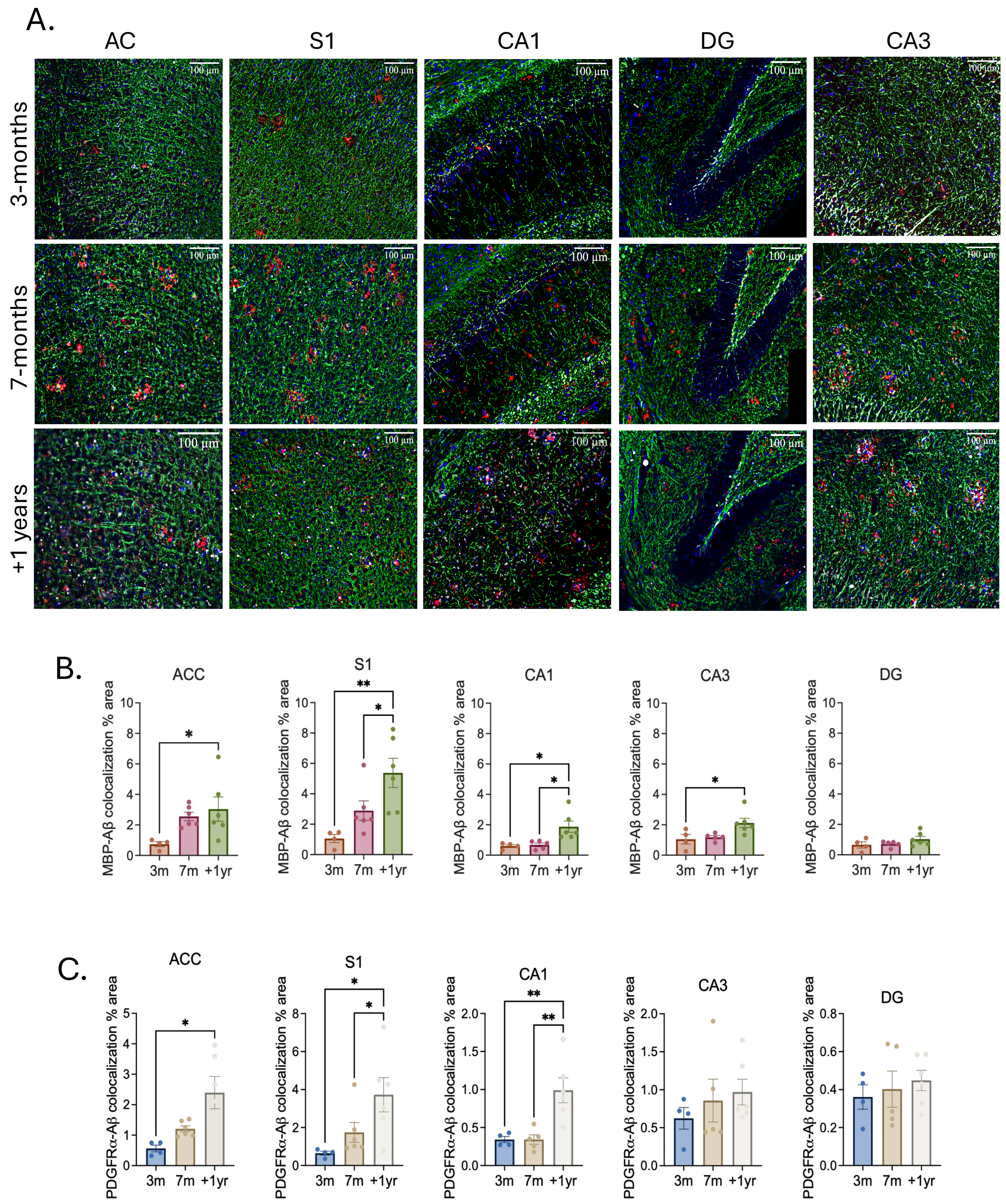

A.
AC
S1
CA1
DG
CA3
3-months
7-months
+1 years
100 µm
B.
ACC
S1
CA1
CA3
DG
MBP-Aβ colocalization % area
3m
7m
+1yr
C.
ACC
S1
CA1
CA3
DG
PDGFRα-Aβ colocalization % area
3m
7m
+1yr

**Supplementary Figure 2. Quantification of MBP and PDGFRα colocalization with Aβ plaques across cortical and hippocampal regions.**

(A) Representative immunofluorescence images of $App^{NL\text{-}G\text{-}F/NL\text{-}G\text{-}F}$ mice at 3 months, 7 months, and +1 year across all ROIs, showing MBP (green), Aβ-plaque (red), PDGFRα (white), and DAPI (blue). Scale bars = 100 µm. (B) MBP–Aβ colocalization (% area) in ACC, S1, CA1, CA3, and DG across disease stages (3 months, 7 months, +1 year) in $App^{NL\text{-}G\text{-}F/NL\text{-}G\text{-}F}$ mice. (C) PDGFRα–Aβ colocalization (% area) across the same regions and timepoints in same $App^{NL\text{-}G\text{-}F/NL\text{-}G\text{-}F}$ mice. Colocalization values represent the proportion of MBP or PDGFRα signal overlapping with Aβ plaques within each ROI. Data are presented as mean ± SEM. Statistical comparisons were performed using one-way ANOVA with Tukey's correction for multiple comparisons. *$p < 0.05$, **$p < 0.01$.
